# Polyploidy of MDA-MB-231 cells drives increased extravasation with enhanced cell-matrix adhesion

**DOI:** 10.1101/2024.06.28.601261

**Authors:** Satomi Hirose, Tatsuya Osaki, Roger D. Kamm

## Abstract

Metastasis, the leading cause of cancer-related deaths, involves a complex cascade of events, including extravasation. Despite extensive research into metastasis, the mechanisms underlying extravasation remain unclear. Molecular targeted therapies have advanced cancer treatment, yet their efficacy is limited, prompting exploration into novel therapeutic targets. Here, we showed the association of polyploidy in MDA-MB-231 breast cancer cells and their extravasation, using microfluidic systems to reproduce the in vivo microvascular environment. We observed enhanced extravasation in polyploid cells alongside upregulated expression of genes involved in cell-substrate adhesion and cell mechanical dynamics. These findings offer insights into the relationship between polyploidy and extravasation, highlighting potential targets for cancer therapy.

## Introduction

Cancer is a leading cause of death worldwide, with metastatic complications contributing to over 90% of cancer-related deaths ^1^. The metastatic cascade involves a sequence of events, including invasion from primary tumors, intravasation, circulation in the blood, extravasation through the vascular walls, and eventual colonization ^2^. While the metastatic process has been widely studied, the mechanisms and biological drivers of extravasation remain unclear.

Over the past three decades, molecular targeted therapies have undergone extensive investigation and refinement, yielding notable clinical advancements in cancer management. These therapies selectively target specific molecular pathways, effectively inhibiting cancer growth, progression, and invasive behavior. However, the current repertoire of molecular-targeted therapies remains limited ^3,4^. Moreover, heterogeneity among cancer cells poses significant challenges in treatment strategies. Different cancer subtypes with varied genetic (e.g., tumor and host mutation) and epigenetic (e.g., methylation and chromatic accessibility) changes may respond differently to treatment, where some patients exhibit significant resistance to existing cancer therapies ^5^. Tumor subtypes also cause variations in cellular mechanical properties ^6^. In the process of extravasation, of the many circulating tumor cells, only a few manage to successfully intravasate from primary tumors and form metastases ^7^. Understanding the specific subtype characteristics that drive cancer cells to progress to form metastases is crucial. Since cancer cells undergo compression and deformation to pass through vessel walls, various mechanical stimuli likely contribute to this process. Identifying these drivers of extravasation holds promise for uncovering novel therapeutic targets, potentially catalyzing breakthroughs in cancer treatment strategies.

MDA-MB-231 is one of the most widely used triple-negative breast cancer cell lines, carrying mutations in the B-Raf proto-oncogene serine/threonine kinase (*BRAF*) and neurofibromin 1 (*NF1*) genes. The histone H2B-enhanced green fluorescent protein (EGFP) fusion protein has been shown to be able to be incorporated into the chromatin of living cells without affecting their cell cycle and viability ^8,9^ and allows high-resolution imaging of chromosomes. While many studies have used H2B-EGFP to label cancer cells, it could be lost in cancer cells due to the methylation of promoters and other genes.

Our research group has been developing microfluidic systems capable of replicating in vivo environments, including intricate 3D microvascular networks. These platforms represent physiologically relevant tissue models, enabling the in vitro observation of cancer cell extravasation dynamics ^10,11^. In experiments studying the extravasation of H2B-EGFP-labeled MDA-MB-231 cells in these devices, we found that cells lacking the H2B-EGFP label exhibited increased extravasation and tended to be polyploid. This observation led us to hypothesize that polyploid cells may demonstrate a heightened propensity for extravasation. While in vivo studies have linked the presence of polyploid giant cancer cells to worse prognoses in certain cancers, we could find no prior published results directly showing the association between polyploidy and extravasation.

We then conducted further experiments and found that polyploid MDA-MB-231 cells exhibit enhanced extravasation within 3D microvascular networks generated in microfluidic devices. Our findings indicate that increased extravasation correlates with the overexpression of integrin subunit beta 1 (*ITGB1*), fibronectin 1 (*FN1*), SH3 and PX domains 2A (*SH3PXD2A*/*TKS5*), vinculin (*VCL*), and AHNAK nucleoprotein (*AHNAK*), which are known to regulate scaffold formation and cell-substrate adhesion. In addition, overexpression of vimentin (*VIM*) and ras homolog family member A (*RHOA*), which are known to regulate cellular mechanical dynamics by upregulating contractility and elasticity, was observed. Collectively, these findings offer a potential explanation for the elevated polyploidy observed in metastatic cancers and provide valuable insights into potential new targets for cancer treatment.

## Results

### EGFP-negative MDA-MB-231 cells extravasate more than EGFP-positive MDA-MB-231 cells

Extravasation was observed in the microvascular networks generated in the microfluidic devices according to the protocol previously established in our group. Briefly, seven days after endothelial cells (ECs) and fibroblasts (FBs) were seeded in a fibrin gel, perfusable microvascular networks were established in the devices by vasculogenesis (**Fig. S1**). These 3D microvascular networks have been used extensively to model physiologically relevant tissues. Extravasation was observed 48 hours after perfusing H2B-EGFP and cytomegalovirus (CMV)-tdTomato tagged (dual-colored) MDA-MB-231 breast cancer cells through these networks (**Fig. 1A**). Counting the extravasated and intravascular cells revealed that extravasated cells accounted for approximately 7% of all cells (**Fig. S2B**). Unexpectedly, half of the extravasated cells did not express EGFP in their nuclei (**Figs. 1B and S2C**), whereas these dual-colored cells should express EGFP in the nuclei and tdTomato in the cytoplasm (**Fig. 1C**). Flow cytometry of the dual-colored cells confirmed that most expressed EGFP but a smaller fraction (∼20%) lacked EGFP expression (**Fig. 1D**). In addition, 6% of the cells lacked tdTomato expression. Since only cells expressing tdTomato were observable within the device, 85% of the perfused cells in the vascular network should be expressing EGFP.

**Fig. 1.**
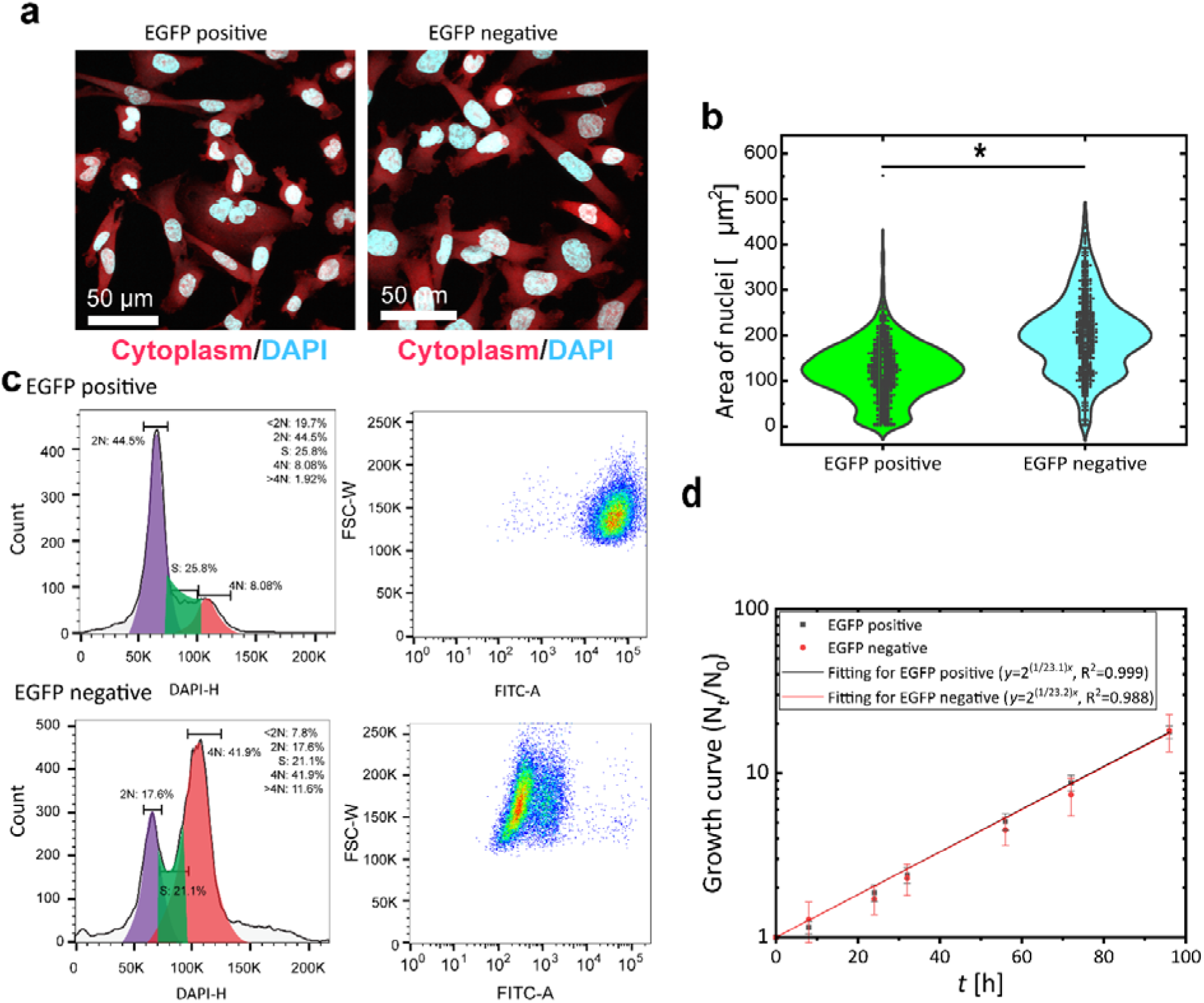
Extravasation of MDA-MB-231-H2B-EGFP-CMV-RFP (double-colored) cells in the microvascular network. (**a**) Representative confocal images of double-colored cells perfused through the microvascular networks. One extravasated cell lacked H2B-EGFP expression. (**b**) The ratio of EGFP negative cells among the extravasated cells in the microvascular networks (*N* = 5, *n* ≥ 19). (**c**) Gene cassette of CMV-H2B-EGFP and CMV-tdTomato were inserted by lenti viurs (**d**) Flow cytometry of dual-colored cells and cell sorting into EGFP-positive and EGFP-negative cells. Comparison of (**e**) counts and (**f**) rate of extravasation of EGFP-positive cells and EGFP-negative cells (*N* = 6, *n* ≥ 18). Error bars show standard deviation. Significant changes were assessed using a *t*-test. Key: *, *p* < 0.01.

This result led to two hypotheses: (1) Insertion of H2B-EGFP affected extravasation, and (2) lacking the EGFP label caused increased extravasation. The first hypothesis was addressed by comparing the extravasation rate with the CMV-tdTomato red fluorescent protein (RFP) tagged (single-colored) MDA-MB-231 cells (**Figs. S2A and S2B**). The dual-colored and single-colored cells (CMV-tdTomato-MDA-MB-231) showed the same level of extravasation. The second hypothesis was addressed by presorting dual-colored cells via fluorescence-activated cell sorting (FACS) into EGFP-positive and EGFP-negative subpopulations. Notably, the extravasation rate was significantly higher for EGFP-negative cells than for EGFP-positive cells, which showed almost the same level of extravasation as the dual-colored cells before sorting (**Figs. 1E and 1F**). These results suggested that the incorporation of H2B-EGFP does not affect extravasation behavior. However, cells that lose their EGFP label show increased extravasation, indicating that the process of losing EGFP labels changes the cellular mechanical dynamics to extravasate.

### EGFP-negative MDA-MB-231 cells tend to be proliferative polyploid cells

We had also noted that the EGFP-negative cells tended to have larger nuclei, suggesting that they might be polyploid cells and may have become stuck in the G2/M phase of the cell cycle. To estimate the cell cycle status of EGFP-positive and -negative cells, nuclei were stained with 4′,6-diamidino-2-phenylindole (DAPI), and nuclear size was determined (**Fig. 2A**). Interestingly, EGFP-negative cells appeared larger than EGFP-positive cells (**Fig. S3**), and the area of their nuclei was significantly larger than that of EGFP-positive cells, as determined by confocal microscopy (**Fig. 2B**). Flow cytometry showed that EGFP-positive cells exhibited a typical pattern in which around half remained in the G0/G1 phase, while the others were in the S or G2/M phases ^12^. In contrast, most EGFP-negative cells contained more DNA than diploid cells (**Fig. 2C**). These results suggest that most EGFP-negative cells are either G2/M arrested or polyploid. Subsequently, their proliferation rate was measured to investigate whether they were G2/M arrested or proliferating as polyploid cells. The proliferation rate of EGFP-negative cells was the same as that of EGFP-positive cells, indicating that EGFP-negative cells are proliferative polyploid cells (**Fig. 2D**).

**Fig. 2.**
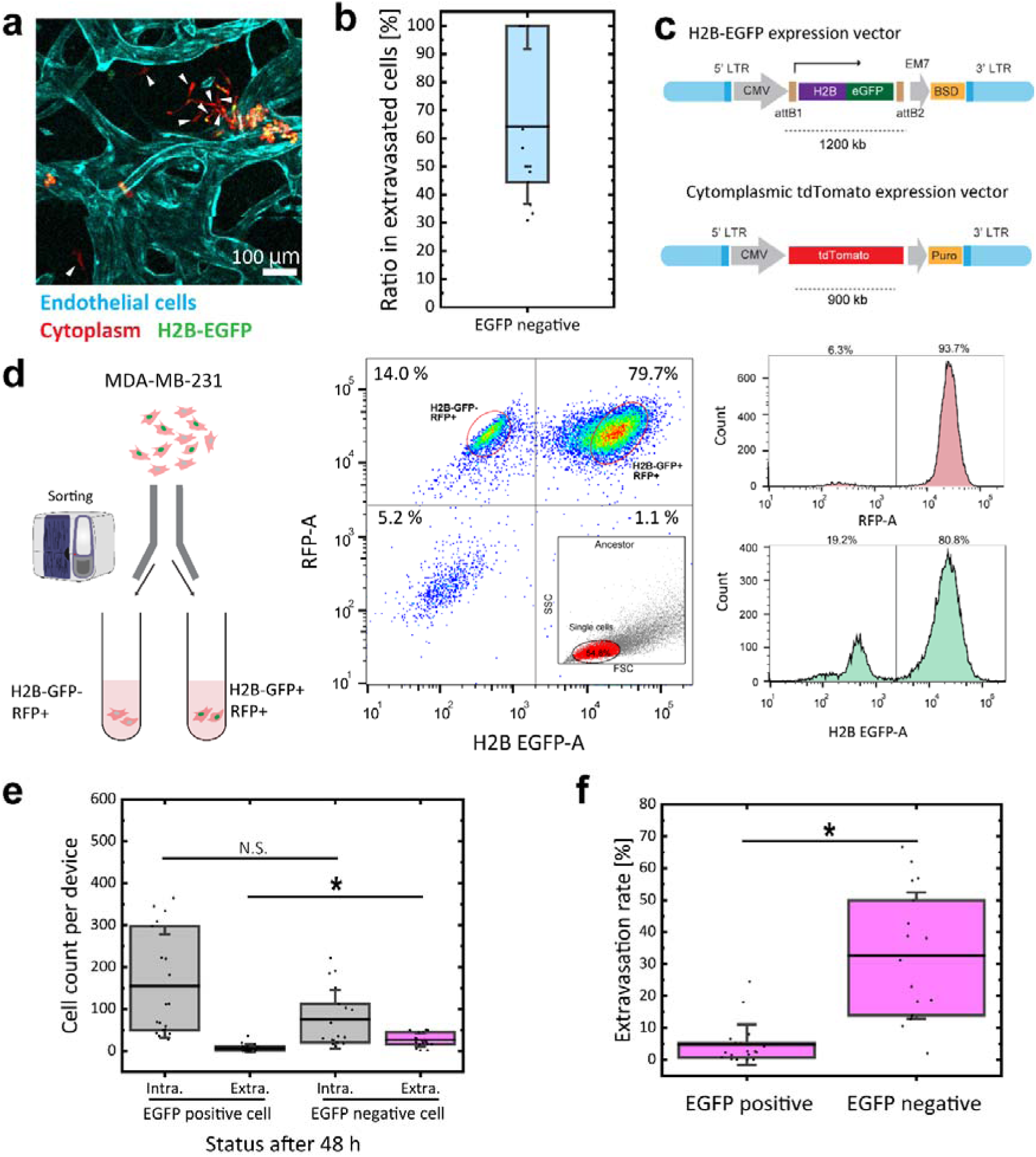
Features of EGFP-positive and EGFP-negative cells. (**a**) Representative images showing the morphology of the sorted cells. (**b**) Comparison of nuclei size between EGFP-positive and EGFP-negative cells (*N* = 10, *n* = 1593 or 1301 cells). Significant changes were assessed using a *t-*test. Key: *, *p* < 0.01. (**c**) Flow cytometry analysis depicting the proportions of sorted cells in each cell cycle phase. (**d**) The proliferation rate of the sorted cells over time (*n* ≥ 10). Error bars show standard deviation.

### Gene upregulation accompanying defective EGFP expression

To further investigate the biological changes occurring with the defect in EGFP expression, bulk next-generation RNA sequencing (RNA-seq) was performed on EGFP-positive and EGFP-negative cells. The volcano plot showed that H2B clustered histone 11 (*H2BC11*) was dramatically downregulated (**Fig. 3A**), indicating that the EGFP label is lost along with the histone H2B. EGFP-negative cells showed little *H2BC11* expression, possibly due to methylation in the promoter regions causing gene silencing, which is common in various tumors ^13^. These partial gene silencing events are a characteristic feature of cancer cells, explaining their heterogeneity ^14^. Conversely, SPANX family member *B1* (*SPANXB1*) expression was significantly higher in EGFP-negative cells than in EGFP-positive cells, indicating that *SPANXB1* is probably involved in the process through which cells lose H2B-EGFP. Gene ontology (GO) enrichment analysis (cellular component) further revealed that the “cell-substrate adhesion,” “extracellular matrix organization,” and “mesenchymal cell differentiation” gene pathways were all activated. Notably, it also indicated strong upregulation of KRAS proto-oncogene GTPase (KRAS) signaling (**Fig. 3F**), which could be related to the polyploidy of cells, as observed above ^15^.

**Fig. 3.**
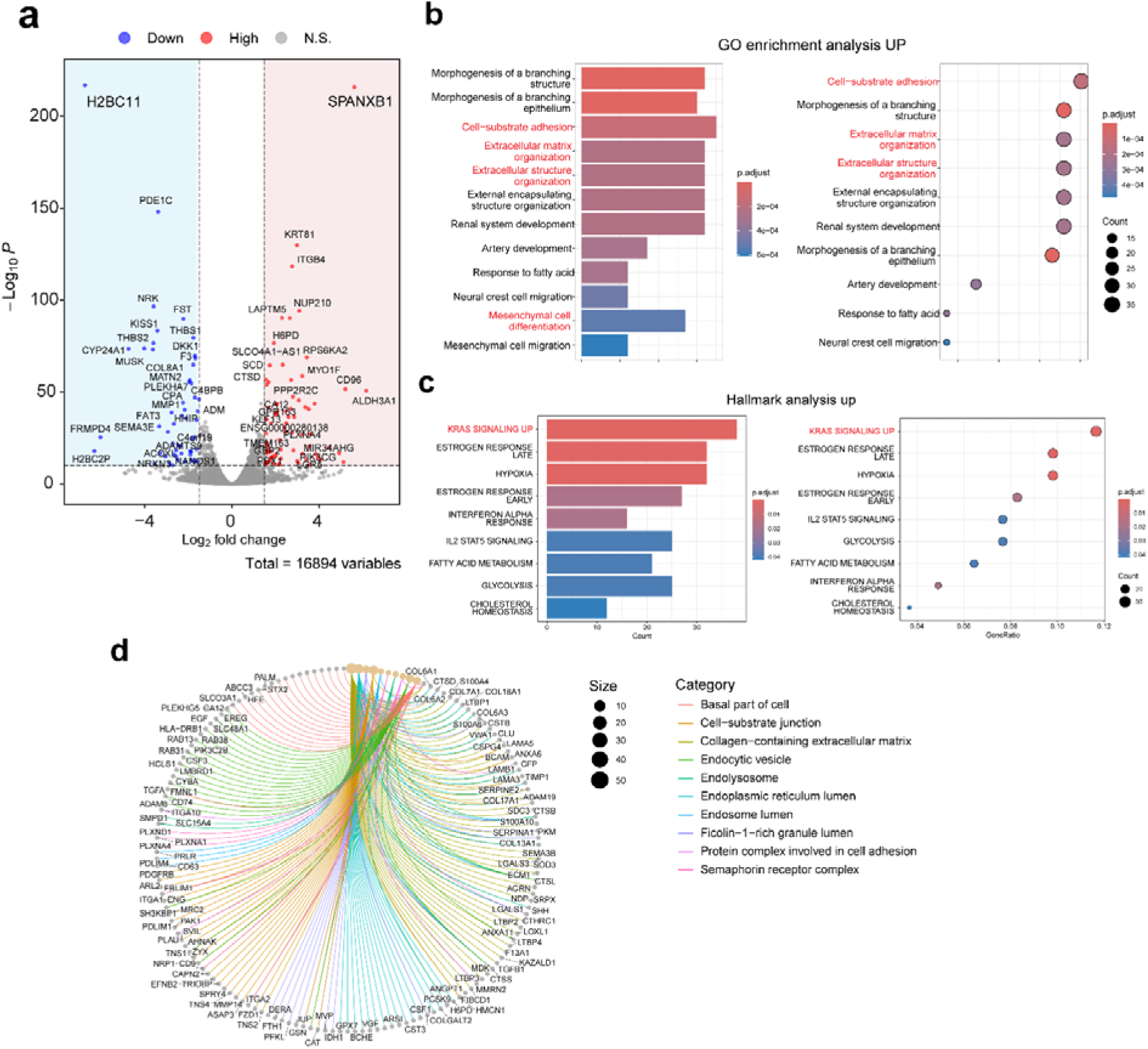
Genes differentially expressed between the EGFP-negative and EGFP-positive subpopulations by RNA-seq. (**a**) Volcano plot illustrating the differential expression of genes between the subpopulations. (**b**) GO enrichment analysis (cellular components) highlighting the pathways upregulated in the EGFP-negative subpopulation. (**c**) Identification of upregulated hallmarks in the EGFP-negative subpopulation. (**d**) Gene concept network plot showing the major pathways associated with the differentially expressed genes.

### Polyploidy in MDA-MB-231 correlates with extravasation

To confirm the association between cell polyploidy and extravasation, dual-colored MDA-MB-231 cells were sorted based on the size of their cell nucleus. After staining cell nuclei with 1 µg mL^−1^ Hoechst 33342, cells with nuclei in the bottom 40% were presorted into the “Small” group, and cells with nuclei in the top 1%–3% were sorted into the “Large” group (**Figs. 4A and 4B**). After sorting and subsequently culturing the cells for 10 days, sub-clones of both populations were perfused into microvascular networks. Consistent with expectations, the Large group showed a higher extravasation rate than the Small group or the dual-colored cells before sorting (**Figs. 4C and 4D**).

**Fig. 4.**
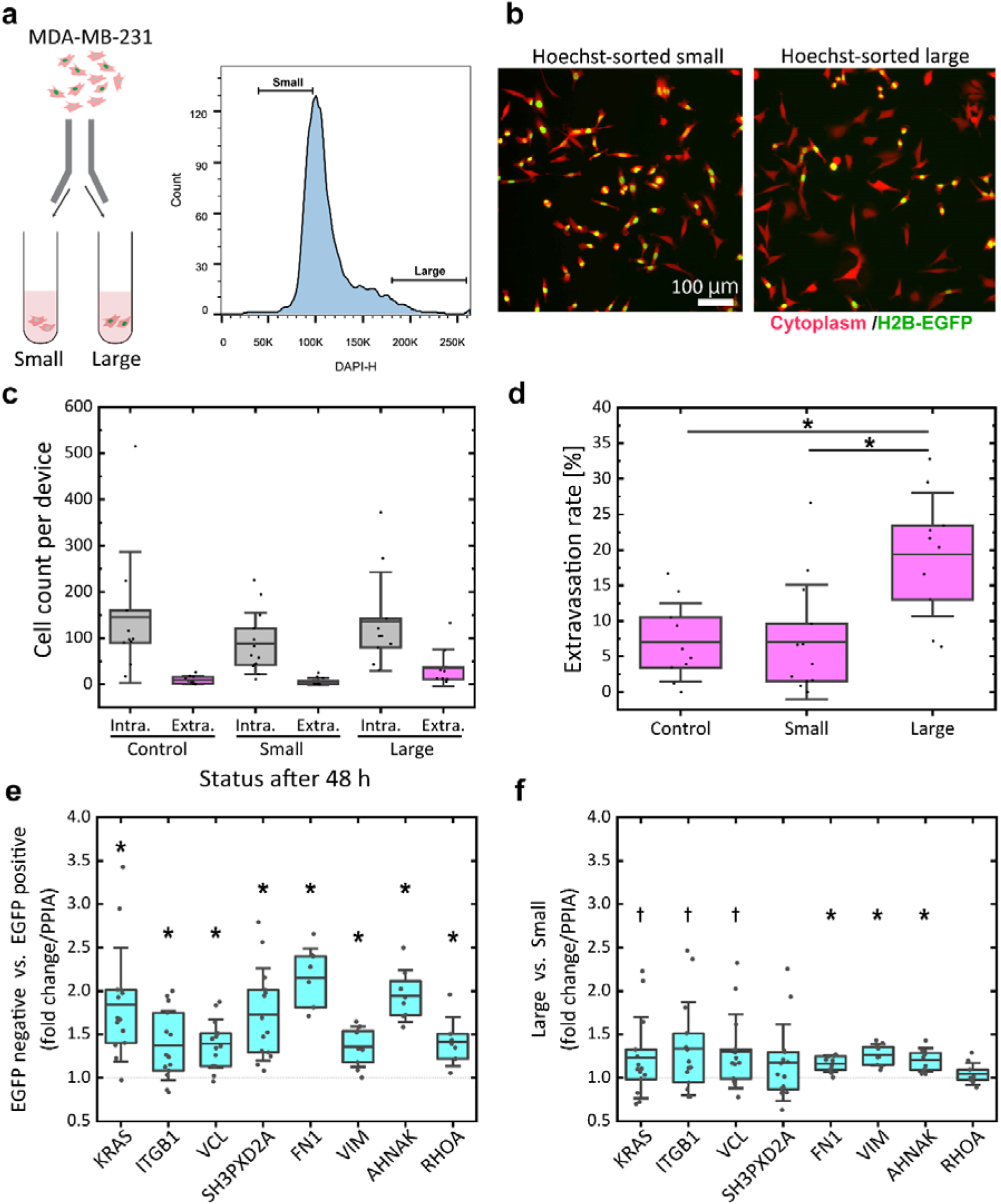
Extravasation rates and gene expression of MDA-MB-231 cells sorted by nucleus size. (**a**) Sorting MDA-MB-231 cells into small and large nuclei groups. (**b**) Representative images showing the morphology of the sorted cells. Comparison of (**c**) counts and (**d**) rate of extravasation in the small and large nuclei groups (*N* = 4, *n* ≥ 10). Comparison of relative gene expression semi-quantified in real-time RT-PCR in (**e**) EGFP-negative versus EGFP-positive cells and (**f**) cells with large versus small nuclei. *PPIA* was used as the housekeeping gene (*N* = 3 or 4, *n* ≥ 7). Error bars show standard deviation. Significant changes were assessed using a *t*-test. Key: *, *p* < 0.01; †, *p* < 0.10.

We also examined the changes in gene expression accompanying the increased extravasation by real-time PCR analyses of genes involved in scaffold formation and cell-substrate adhesion (*ITGB1*, *FN1*, *VCL*, *SH3PXD2A* [*TSK5*], and *AHNAK*) and cytoskeletal mechanical stiffness (*VIM* and *RHOA*) (**Fig. 4E**). The expression of all these genes was higher in the Large group than in the Small group. However, the differences tended to be smaller than those between EGFP-negative and EGFP-positive cells, consistent with the smaller difference in extravasation rate between the Large and Small groups than between the EGFP-negative and EGFP-positive cells.

Flow cytometry of cells cultured for the same duration as cells used for perfusion indicated that the Small and Large groups contained 56% and 81% G2/M or polyploid cells, respectively (**Figs. S3 and S4**), while the extravasation rate was ∼20% in the Large group compared to ∼45% for the EGFP-negative cells (**Fig. 1F**). This finding indicates that despite the higher proportion of polyploid cells in both the Large and Small groups compared to EGFP-negative cells, their extravasation rates were lower. This discrepancy could potentially be attributed to the effects of Hoechst 33342 staining on live cells. While Hoechst 33342 is generally considered less toxic than some other stains and even used for flow-sorting of boar spermatozoa ^16^ and murine hematopoietic stem cells ^17^, since Hoechst stains cell nuclei as a DNA-binding ligand, it has toxicity especially at a higher concentration since it inhibits DNA synthesis ^18,19^. Our study used Hoechst33342 at a concentration less than 20% of the recommended concentration ^20^, and the fluorescence dissipated within a day. However, the observation that most cells became polyploid, even in the group expected to be diploid or have lower DNA content, suggests that Hoechst staining and subsequent sorting may have influenced the cell cycle. The doubling time of the cells cultured for the same duration as cells used for perfusion was 22.7 hours for the Large group, consistent with the cells before sorting, and 26.3 hours for the Small group (**Fig. S4**). Considering that at the time of sorting, most cells sorted into the Small group should be in the G0 or early G1 phase, it is plausible that the Small group was affected by DNA synthesis inhibition with Hoechst more than the Large group. Despite these considerations, we still considered the comparisons between the Small and Large groups with differing polyploidy populations to be informative as an EGFP-independent comparison between groups.

## Discussion

Despite the critical role of cancer cell extravasation in metastasis, its underlying mechanisms and molecular factors remain inadequately studied due partly to their heterogeneity. Here, we used a 3D microvascular network within microfluidic devices to perfuse MDA-MB-231 breast cancer cells and observe their extravasation, aiming to elucidate the features of the cell group exhibiting heightened extravasation. Our findings revealed that the subset exhibiting higher extravasation rates contained a higher proportion of polyploid cells, which also exhibited increased expression of genes involved in cellular mechanical dynamics. Interestingly, in a serendipitous discovery, while observing extravasation in vitro using MDA-MB-231 cells harboring the H2B-EGFP insertion, we observed a subpopulation that repressed histone H2B and associated EGFP expression, exhibiting enhanced extravasation compared to other cells (**Fig. 1**). The H2B-EGFP fusion protein, known to be properly incorporated into chromatin without affecting the cell cycle ^8^, is commonly used for cell nuclei and chromatin imaging ^21^. To date, the effects of H2B-EGFP insertion or deletion on cells have not been reported. Given that gene silencing caused by promoter methylation is common in cancer cells exhibiting high heterogeneity ^13^, the observed subpopulation of EGFP-negative cells likely arises naturally due to heterogeneity. This observation also serves as a cautionary note regarding the sorting of cancer cells, as it could be unexpectedly selective under certain labeling due to their high heterogeneity.

The EGFP-negative cells were found to have more DNA in their nuclei despite exhibiting an unaltered proliferation rate, consistent with proliferative cancers, which are highly proliferative and also tend to favor polyploid conditions (**Fig. 2**). In cancer cells, polyploidy is usually associated with cellular senescence, and polyploid cells are considered to have a slow or arrested cell cycle ^22,23^. However, since cancer cells could lack tumor suppressor genes that regulate cell proliferation and cytokinesis ^24,25^, the association between polyploidy and senescence in cancer cells is unclear, and the existence of proliferative polyploids is unsurprising. Indeed, it was reported in a mouse model that cells that overcame DNA replication stress to drive unlimited proliferation in culture exhibited higher polyploidy ^26^. In addition, it is notable that 90% of human solid cancers are aneuploid, with 20% being polyploid ^27^. Moreover, we showed that the *KRAS* signaling pathway was upregulated in EGFP-negative cells, according to the hallmark analysis (**Fig. 3E**), which could be associated with their polyploidy. The KRAS pathway is considered the most common oncogenic driver in human cancers and plays an important role in various cancer behaviors, including progression, survival, and migration ^28^. *KRAS* is generally expressed in breast cancer cells, including MDA-MB-231 ^29^. Some studies have observed that upregulation or mutation of *KRAS* is associated with polyploidy ^15,30^.

Among the factors possibly contributing to the increased extravasation rate of EGFP-negative cells, KRAS is particularly interesting, given its known propensity to promote the migration and invasion of MDA-MB-231 cells via the upregulation of *VIM* and *FN1* ^31^. Moreover, the GO enrichment analysis indicated the upregulation of “cell-substrate adhesion,” with the upregulated expression of *ITGB1*, *FN1*, *VCL*, *SH3PXD2A*, and *AHNAK* confirmed by real-time PCR. The GO enrichment analysis also indicated the upregulation of “mesenchymal cell differentiation,” with the upregulated expression of *VIM* and *RHOA* confirmed by real-time PCR. The increased extravasation rate and increased expression of these genes in highly polyploid cells were also reproduced by comparison of cell populations sorted by nuclear size independent of EGFP expression **(Fig. 4)**.

A previous study from our group demonstrated that ITGB1 is required for tumor cell invasion past the basement membrane following the clearance of the endothelial barrier during the extravasation process ^32^. It was proposed that ITGB1-rich extracellular adhesions maintain protrusions past the endothelium, allowing focal adhesion proteins and F-actin to be recruited to the tips of these protrusions and resulting in transmigration via actomyosin contraction at the protruding edge. These protrusions of extravasated cells were marked by colocalized punctate regions of F-actin and activated ITGB1, VCL, and TKS5 at the tips. The upregulation of *ITGB1*, *VCL*, and *TKS5* observed in EGFP-negative cells, accompanied by increased extravasation, is consistent with our previous findings.

In addition, in our study, we observed decreased extravasation of EGFP-negative cells caused by the supplement of 0.5 mM RGDS peptide, which blocks the adhesion of FN1 and integrins **(Fig. S5)**. Notably, we also observed upregulated *FN1* expression in the subpopulations of EGFP-negative and large-nuclei cells **(Fig. 4)**. However, it remains unclear if the secretion of FN1 from tumor cells supports extravasation by strengthening focal adhesions since the extracellular matrix is originally rich in FN1 secreted by ECs and FBs. These results suggest that the increased extravasation observed in polyploid cells is due to the formation of more protrusions past the endothelium, leading to complete extravasation.

AHNAK is also an important scaffold protein, and while its precise function remains unclear, it has been implicated in diverse biological processes ^33^. In cancer cells, AHNAK is reported to be essential for migration and invasion with actin-dependent pseudopod protrusion ^34^ and is highly expressed in some tumors, including metastatic human breast cancer ^35,36^. Considering that the formation of actomyosin-rich protrusions is required for extravasation ^32,37^, the upregulated expression of *AHNAK* might enhance the stability of the scaffold and actomyosin during extravasation, contributing to its efficiency. A previous study from our group has also shown that adding cytochalasin D, which disrupts actin polymerization and AHNAK distribution, significantly decreased the extravasation of MDA-MB-231 cells ^10^. The associated regulation of elasticity and contractility in polyploid cells might contribute to higher extravasation rates since cancer cells are known to optimize these properties during migration for increased efficiency ^38^.

VIM is widely accepted as a major migration enhancer ^39^ and is normally highly expressed in MDA-MB-231 since it is one of the major epithelial-mesenchymal transition markers ^40,41^. Studies have shown that *VIM* expression is elevated in drug-induced polyploid MDA-MB-231 cells, accompanied by greater migration in wound healing assays ^42^ and directional migration ^43^. Notably, in vivo data on human patients with breast cancer have shown higher *VIM* expression in metastatic tumors, followed by primary tumors with metastasis, compared to primary tumors without metastasis and benign tumors ^42^. These reports are consistent with the changes in *VIM* expression in polyploid cells observed in our study. VIM is also known to enhance cell stiffness, increase the resistance to deformation and compressive strain, and maintain cellular integrity ^44,45^, which could be a key factor in executing extravasation and survival afterward.

The fact that VIM increases cell stiffness while increasing extravasation of polyploid cells with large cell nuclei and cell volume may seem contradictory, considering the physical barrier to passage through the blood vessel wall. However, previous studies using MDA-MB-231 *VIM* knockdown cells have also shown that VIM is required for invasion in transwell assays ^46^. Additionally, we observed decreased extravasation of EGFP-negative cells following the inhibition of VIM by a 48-hour pre-treatment with 5 mM metformin ^47,48^ **(Fig. S5)**.

RHOA is upregulated in various human cancers ^49^ and is considered to contribute to invasion ^50^ by regulating the contractility of actomyosin ^51^. Previous studies have shown that knocking down *RHOA* ^37^ and inhibiting the function of myosin heavy chain 2 (MYH2) and actomyosin contractility with blebbistatin ^10^ decreased the extravasation efficiency of MDA-MB-231 cells. For cancer cell invasion, it has been reported that mesenchymal migration can transition into an amoeboid type ^52^, another mutually interchangeable form of migration ^53^, when cells enter confined 3D environments. Therefore, investigating gene expression in 3D cultures and confined environments, such as in situ observation of extravasating cells, would provide further insights.

Altogether, these findings indicate that polyploidy in MDA-MB-231 cells increases successful extravasation through enhanced scaffold formation, improved cell-substrate adhesion, and changes in cytoskeletal and cellular mechanics. Polyploidy is considered an important feature of cancer determining its prognosis, as many clinical studies have reported that higher polyploidy may result in poor prognosis ^54^. The number of polyploid giant cancer cells was higher in metastatic than in primary tumors of patients with breast cancer ^42^. However, the mechanism remains unclear, but it is believed to mainly be due to polyploid cells’ therapeutic resistance ^55,56^ and higher dormancy and reactivation ^57,58^. In addition to these factors, our study shows that high extravasation rates of polyploid cells may contribute to poor prognosis. The changes in gene expression involved in the cellular mechanical dynamics of cancer cells observed in our study may also contribute to other stages in the metastatic cascade, such as invasion and resistance to damage due to fluid shear stress in the blood circulation. Therefore, our findings might help to explain the elevated polyploidy observed in metastatic cancers and provide valuable insights into potential new targets for cancer treatment.

Notably, contrary to previous studies that investigated the in vitro dynamics of polyploid cells created by chemical reagent or anticancer drug treatment, our study focused on naturally occurring polyploid cells arising from the heterogeneity of cancer cells. While our results align with many findings from previous studies on artificially induced polyploid cells, there may be disparities in their characteristics. Additionally, clinically observed polyploidy in tumors is likely both drug-induced and naturally occurring, and the similarities and differences between their features or prognoses remain unclear. Clarifying these details could offer insights into more personalized targets for cancer treatment.

Finally, the detailed mechanism(s) by which the subpopulation of EGFP-negative cells lose H2BC11 remains unclear. The *SPANXB1* gene, which is exclusively expressed in EGFP-negative cells, is likely related to this process. SPANXB1 has been reported to promote metastasis in triple-negative breast cancer ^59^. Additionally, a previous study indicated that its expression was higher in brain metastatic MDA-MB-231 cells than in the parental MDA-MB-231 cells. In contrast, its expression was similar in tumor and normal tissues from patients with breast cancer ^60^. These findings align with the observations in our study, where *SPANXB1* expression was only detected in the subpopulation exhibiting high extravasation. However, few studies have explored this gene extensively, and there are no clear clues to decipher its causal relationship. Numerous other factors beyond those discussed here may contribute to extravasation in polyploid cells. For example, changes in nuclear volume might well influence nuclear stiffness, thereby affecting the ability of the tumor cell to pass through the endothelial barrier. It is anticipated that as our understanding of gene modulation and its roles becomes clearer in the future, the mechanism by which polyploidy is related to the extravasation of cancer cells will also become clearer.

## Methods

### Cell culture

The following cells were used: immortalized human umbilical vein ECs ^61^, which proliferate indefinitely and allow for more reproducible results; normal human lung FBs (CC-2512; Lonza); and MDA-MB-231 triple-negative breast cancer cells (CRM-HTB-26, ATCC) transfected to express cytoplasmic tdTomato (MDA-MB-231-CMV-RFP/single colored). MDA-MB-231-CMV-RFP cells were further tagged with H2B-EGFP (MDA-MB-231-H2B-EGFP-CMV-RFP/dual colored). MDA-MB-231-H2B-EGFP-CMV-RFP cells were sorted using FACS into EGFP-positive and EGFP-negative subpopulations. Briefly, lentivirus transfer plasmid was constructed from (pLenti6/V5-DEST™ Gateway™ Vector, Thermo Fisher Scientific) as backbone plasmid and entry plasmid (pENTR-H2B-EGFP) by gateway cloning with Manufacturers’ protocol. Then, lentivirus was packaged with pLP.1 (gag/pol), pLP.2 (rev), and pLP/VSVg (envelop as 3rd generation packaging system with ViralPower lentivirus expression system (Thermo Fisher Scientific) in 293FT cells. After lentivirus infection in MDA-MB-231-CMV-RFP with polybrene, the cells were selected using Blasticidin and FACS to obtain MDA-MB-231-H2B-EGFP-CMV-RFP cells (tdTomato and EGFP positive cells). Furthermore, to identify subpopulations after many passages, MDA-MB-231-H2B-EGFP-CMV-RFP cells were further sorted using FACS once again into EGFP-positive and EGFP-negative subpopulations.

The cell lines were cultured at 37°C with 5% CO_2_ in atmosphere. ECs were cultured in VascuLife culture medium (LL-0003; Lifeline cell technology), FBs were cultured in FibroLife VascuLife culture medium (LL-0011; Lifeline cell technology), and MDA-MB-231 cells were cultured in Dulbecco’s modified Eagle’s medium (11995065; Gibco) supplemented with 10% fetal bovine serum (FBS; 26140079; Gibco), 100 U mL^−1^ penicillin and 100 µg mL^−1^ streptomycin (P4333; Sigma-Aldrich). FACS-sorted cells were first expanded, then simultaneously frozen in a sufficient number of vials, and used for experiments after thawing individually.

### Microvascular network formation

The microfluidic devices used in this study to generate microvascular networks were fabricated from polydimethylsiloxane (Sylgard 184; Dow Corning) by standard soft lithography with a central gel channel for cell seeding and two adjacent media channels, as previously described ^62^. The dimensions of the central gel channel were 3 mm (width) × 5 mm (length) × 0.25 mm (height). For seeding into gels, ECs and FBs were harvested from cell culture flasks after five days of culturing post-thawing using ACCUTASE (07922; STEMCELL Technologies) and resuspended in VascuLife supplemented with thrombin (T4648; Sigma-Aldrich). Then, the cell suspension was mixed with the same volume of phosphate-buffered saline (PBS) containing fibrinogen (341578; Sigma-Aldrich) and injected into the gel channel of microfluidic devices. The final concentrations of ECs, FBs, thrombin, and fibrinogen were 3 × 10^6^ cells mL^−1^, 1.5 × 10^6^ cells mL^−1^, 2 U mL^−1^, and 3 mg mL^−1^, respectively. After the gel was polymerized in an incubator at 37°C, VascuLife was injected into both media channels. The VascuLife in the media channels was changed daily, and the ECs formed perfusable vascular networks by vasculogenesis seven days after seeding.

### Quantification of extravasation

MDA-MB-231 cells were harvested from cell culture flasks after five days of culturing post-thawing using 2 mM EDTA (15575020; Thermo Fisher Scientific) in PBS. The cells were resuspended in VascuLife containing 10% FBS at a density of 2 × 10^6^ cells mL^−1^. After aspirating the medium in the media channels of the microfluidic devices, 20 µL of cell suspension was injected into a media channel on one side, followed by 200 µL of Vasculife containing 10% FBS. Due to the head difference, the medium gently flowed to the other side, and the cells were dispersed within the microvascular network. The VascuLife containing 10% FBS in the media channels was changed after 24 hours. The microvascular network was imaged after 48 hours using a confocal microscope (FV1000, Olympus) with a 10× objective lens. The microvascular network in the entire gel channel was imaged by tiling, and images on the horizontal plane (*xy*-plane) were taken at 100 µm intervals in the vertical (*z*-axis) plane. For each device, a 2 mm (width) × 3 mm (length) region of interest was set, avoiding the area near the media channels, and the number of all MDA-MB-231 cells within it was counted manually using the ImageJ software (US National Institutes of Health) with its Cell Counter plugin. When cells clustered or overlapped, the number of cells was estimated to the best of our ability based on cluster size. Cells in the process of extravasation were counted as extravasated when at least half of the cell area was outside the vessels. At least three independent experiments with at least eight devices were performed for each condition. Significant differences between the three conditions were evaluated using Bonferroni’s multiple comparison test. Significant differences between pairs of conditions were evaluated using a *t*-test. A *p* < 0.01 was considered statistically significant.

### Quantification of nuclear size

EGFP-positive and EGFP-negative subpopulations of dual-colored MDA-MBA-231 cells were placed in φ35 glass bottom dishes, fixed with 4% paraformaldehyde (157-8-100, Electron Microscopy Science) in PBS, and stained with 5 µg mL^−1^ DAPI (D1306; Invitrogen). Next, ten microscopic images were captured from two dishes per subpopulation using an epi-fluorescence microscope (Eclipse Ti-S; Nikon) with a 10× lens. Within each image (1392 × 1040 pixels, 0.93 × 0.69 mm), the size of the nuclei of all cells was quantified using ImageJ software. Significant differences between the two conditions were evaluated using a *t*-test. A *p* < 0.01 was considered statistically significant.

### Quantification of proliferation rate

The EGFP-positive and EGFP-negative subpopulations of dual-colored MDA-MBA-231 cells were seeded in a 24-well plate at a density of 1000 cells per well. The next day was defined as 0 hours, and one image of each well was taken every few hours up to 96 hours using the epi-fluorescence microscope with a 4× lens. The medium was replaced every two days. The number of cells in each image (1392 × 1040 pixels, 2.25 × 1.68 mm) was counted manually using the ImageJ software with its Cell Counter plugin, and the doubling time was calculated by fitting the *y* = 2^(1/*a*)*x*^ formula for a plot of elapsed time and cell number.

### Flow cytometry and image-enhanced flow cytometry

To sort the EGFP-positive or EGFP-negative subpopulations, MDA-MB-231 cells were dissociated by treatment with ACCUTASE for 2 min. Next, the cell suspension was filtered with a 70-µm cell strainer (BD Falcon) in 1% BSA in PBS. Then, the cell population was gated by the forward scatter (FSC) and side scatter (SSC) to remove duplet/triplet and then further gated and sored as EGFP-positive and EGFP-negative cells using a BD FACSMelody or FACSAria cell sorter (**Fig. S4**). For cell cycle analysis and sorting of MDA-MB-231 cells with large or small nuclei, the cells were incubated in culture medium containing Hoechst 33342 for 15 min at 37°C and then washed thrice with PBS. Then, the cells were collected, and the same protocols as above were followed for sorting or cell cycle analysis.

Imaging-enhanced flow cytometry (Attune CytPix, Thermo Fisher Scientific) was used to characterize the heterogeneity in the size and morphology of the subpopulations of MDA-MB-231 cells. The cell subpopulations were gated by FSC and SSC and then imaged. The acquired images were then used to estimate the cells’ size, perimeter, and circularity in CytPix, and integrated data (images, FSC, and SSC) were further analyzed in MATLAB (Mathworks). All other analyses and visualization of flow cytometry data were performed using FlowJo software.

### RNA-seq

The EGFP-positive and EGFP-negative subpopulations of MDA-MB-231 cells were lysed using RNeasy Plus Mini Kit (74134; QIAGEN) and kept at −80°C until needed. Next, PolyA selection and stranded RNA-seq library preparation were performed with the NEBNext Ultra II Directional RNA Library Prep Kit for Illumina (E7760; New England Biolabs). Then, bulk RNA-seq was performed by the Genomics Core of the Koch Institute for Integrative Cancer Research (MA, USA) using a 100M paired-end configuration on the Singular G4 platform. The raw reads were demultiplexed based on the tag and mapped to build GRCh37/hg19 of the human reference genome. Next, the mapped reads were quantified using the featureCount in Subread. Differential expression was detected using a Wilcoxon rank sum test and visualized as a volcano plot using R (version 4.3.2) with edgeR package (version 4.0.14), with *p*-values adjusted for multiple testing using the Benjamini–Hochberg method. These clusters were further characterized using bulk hallmark pathway analysis conducted using the enricher function in the R clusterProfiler package (version 4.10.1) with the hallmark and GO gene sets from MSigDB ^63,64^. All count files were deposited into Gene Expression Omnibus (TBA).

### Real-time RT-PCR

Real-time RT-PCR was conducted to semi-quantify the mRNA levels of genes whose expression was elevated in the single-colored, dual-colored, and sorted MDA-MB-231 cells in the RNA-seq data. Total RNA was extracted from each cell group cultured on cell culture dishes using the RNeasy Plus Mini Kit (74134; Qiagen) and converted to cDNA using the Verso cDNA Synthesis Kit (AB1453A; Thermo Fisher Scientific). Following the product manual, the RT-PCR reaction was performed in 20 μL of reaction buffer containing 10 μL of TB green Premix Ex Taq II (RR82WR; Takara), 1 μL of 10 μM forward primer, 1 μL of 10 μM reverse primer, 1 μL of 5 µg mL^−1^ cDNA, and 7 μL of water. The primers used in this study were outsourced as custom DNA oligonucleotides (Integrated DNA Technologies), and their sequences are provided in Table 1. Real-time PCR was performed using a Quantstudio 3 Real-Time PCR System (Thermo Fisher Scientific). Three independent experiments were performed, each with three or four samples for each cell group for each condition. Relative gene expression was calculated using the 2^−ΔΔCt^ method, with the mRNA levels of the target genes normalized to those of peptidylprolyl isomerase A (*PPIA*) as a housekeeping gene. Significant differences between two conditions were evaluated using paired *t*-tests. Significant differences between three or more conditions were evaluated by one-way analysis of variance followed by post-hoc Tukey’s multiple comparison tests. A *p* < 0.01 was considered statistically significant in each test, and a *p* < 0.10 was considered marginally significant.

## Acknowledgments

The authors wish to acknowledge research support from the National Cancer Institute (U54 CA261694). We thank the Koch Institute’s Robert A. Swanson (1969) Biotechnology Center for technical support, specifically flowcytometry core. We also thank Dr. Peter Friedl, Dr. Arhadana Singh and Dr. Carolina Jorgez to prepare the lentivirus construct and obtain stable EGFP and tdTomato expression MDA-MB-231. TO was supported by NIH grant R01MH085802. SH was supported by a JSPS Overseas Research Fellowship.

## Author contributions

SH, TO, and RDK conceived and designed the experiments. SH performed cell experiments and PCR, and analyzed data. TO performed flow cytometry and PCR and analyzed RNA-seq data. SH, TO, and RDK wrote the manuscript.

## Data availability

The RNA-seq data were deposited in the Gene Expression Omnibus database (TBA). All raw data used to generate the graphs in this article can be found at TBA.

## Supplemental materials

**Fig. S1:**
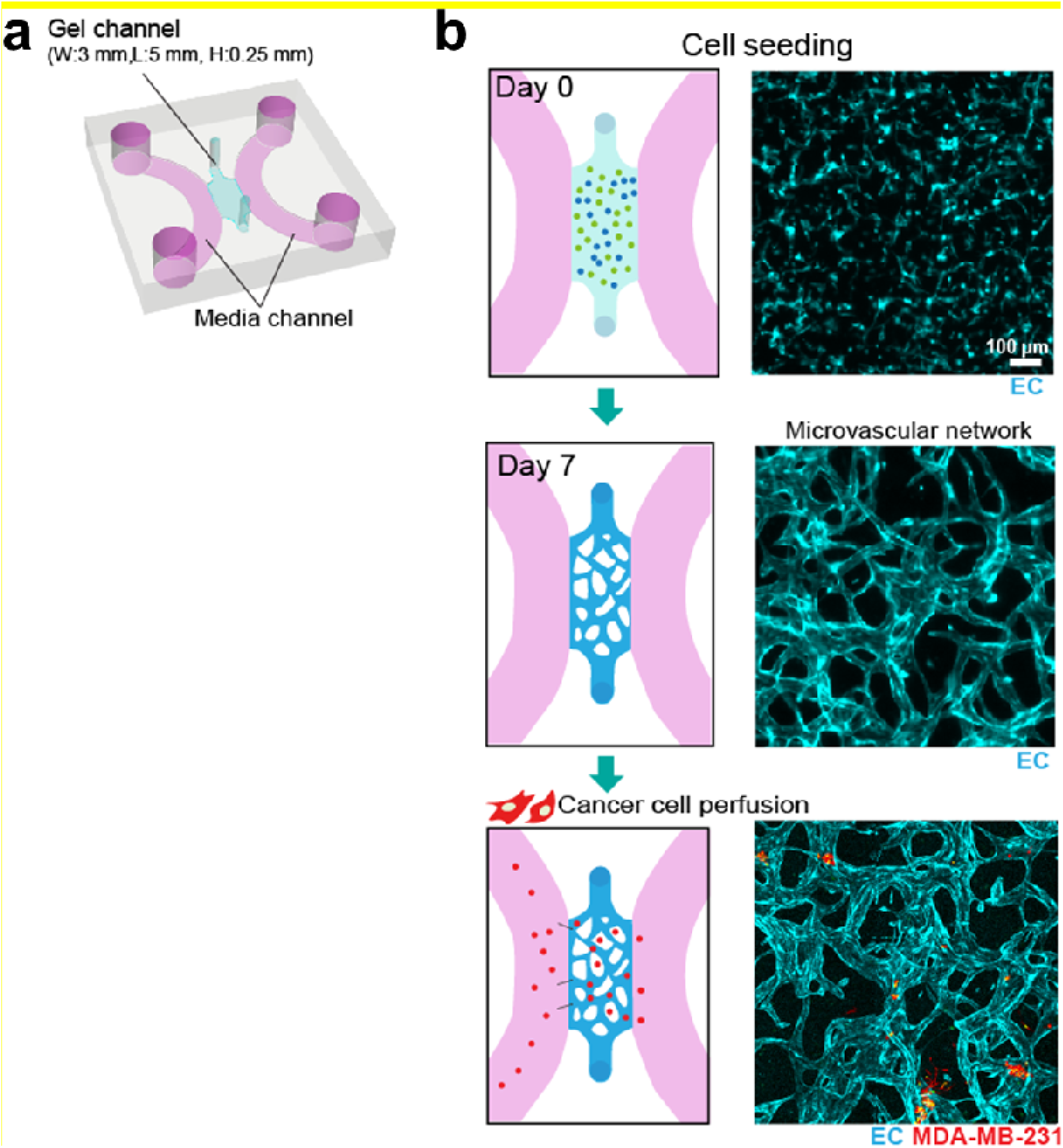
Experimental procedure. (a) Schematic of the microfluidic device. (b) Microvascular network generated in the gel channel.

**Fig. S2:**
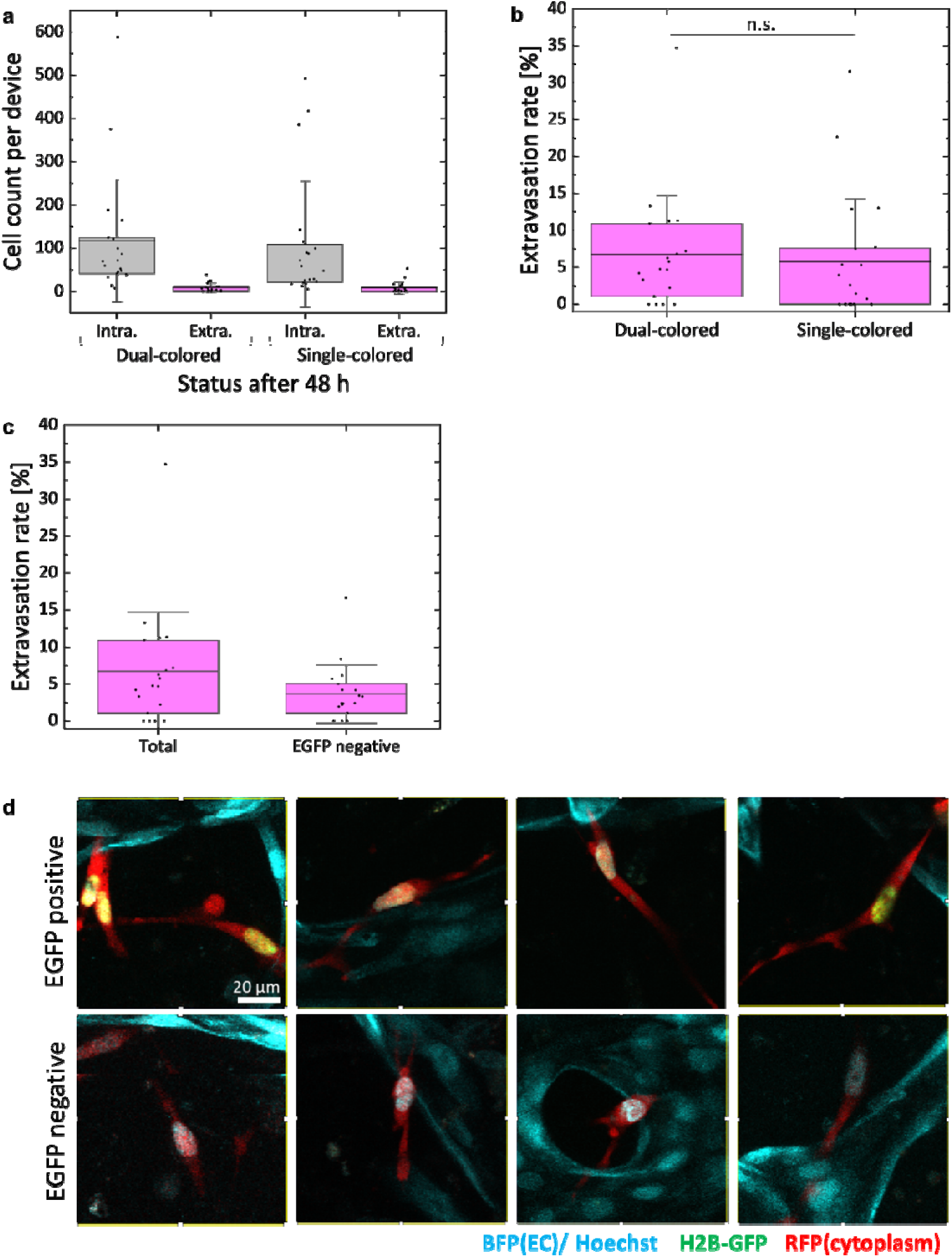
Comparison of (a)counts and (b)rate of extravasation of dual-colored cells (MDA-MB-231-H2B-EGFP-CMV-RFP) and single-colored cells (MDA-MB-231-CMV-RFP). (c) For the results with double colored cells, the number of EGFP negative cells among the extravasated cells in the microvascular networks was counted (*N* = 5, *n* ≥ 19). Error bars show standard deviation. Significant changes were assessed by *t*-test. (d) Representative image of extravasated cells in microfluidic devices.

**Fig S3:**
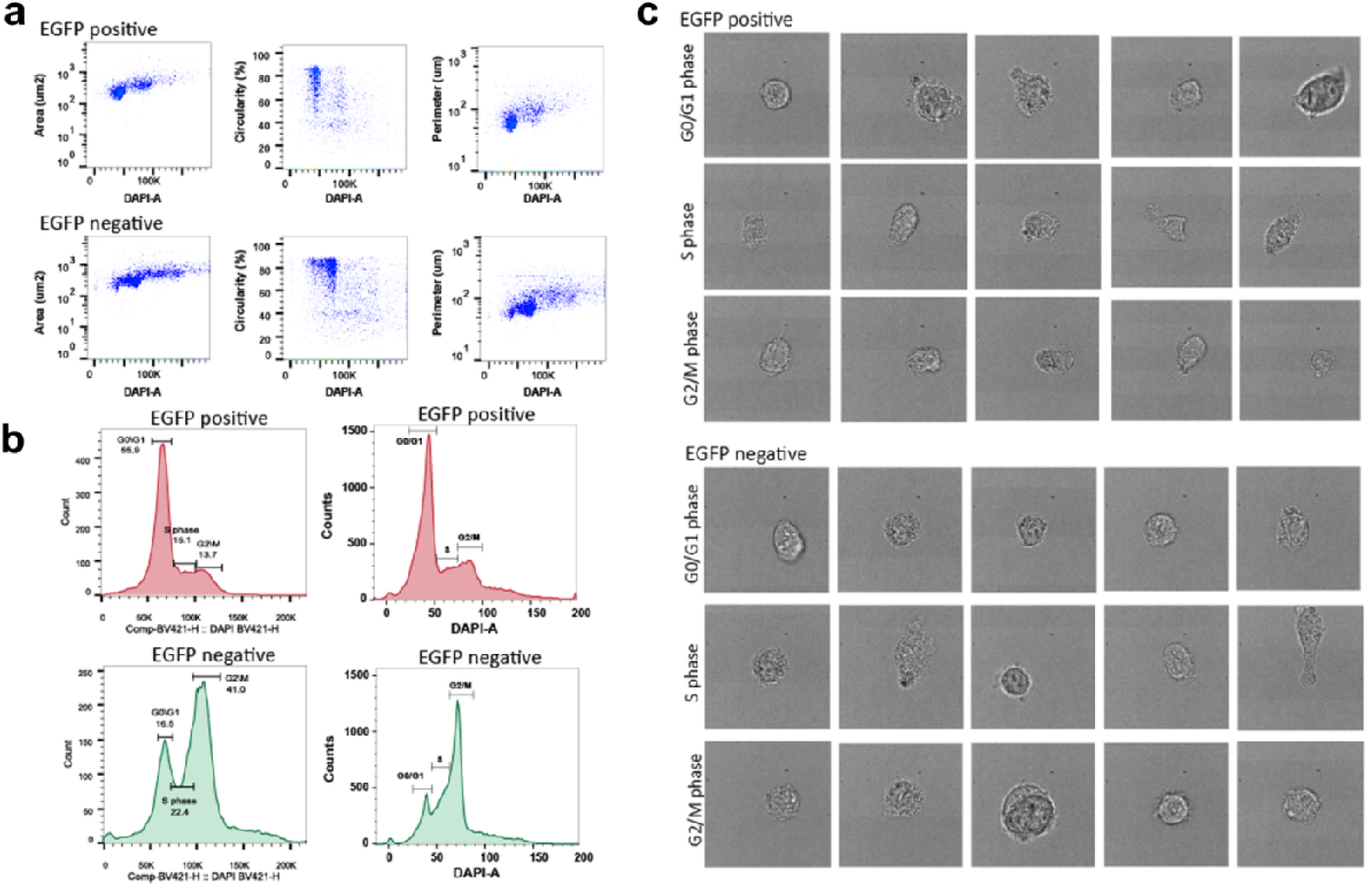
Comparison of the morphology of EGFP positive cell and EGFP negative cells. (a) Distribution of indicators of cell morphology. (b) Cell cycle analysis by using BD FACSMelody (right) or FACSAria (left). (c) Representative image of floating cell morphology.

**Fig. S4:**
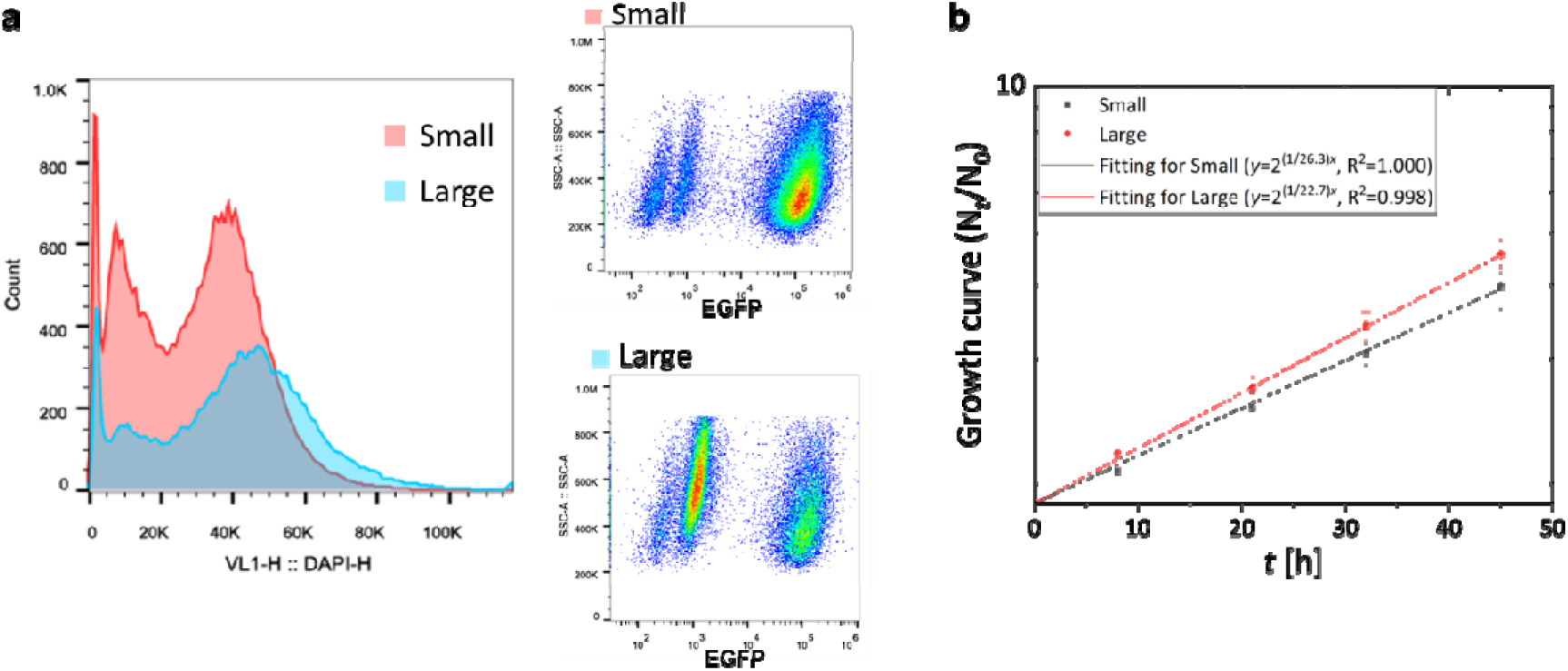
Characterization of cell after cell-sorting with Hoechst staining. (a) Result of flow cytometry. (b) The proliferation rate of sorted cells over time (*n* ≥ 10). Error bars show standard deviation.

**Fig. S5:**
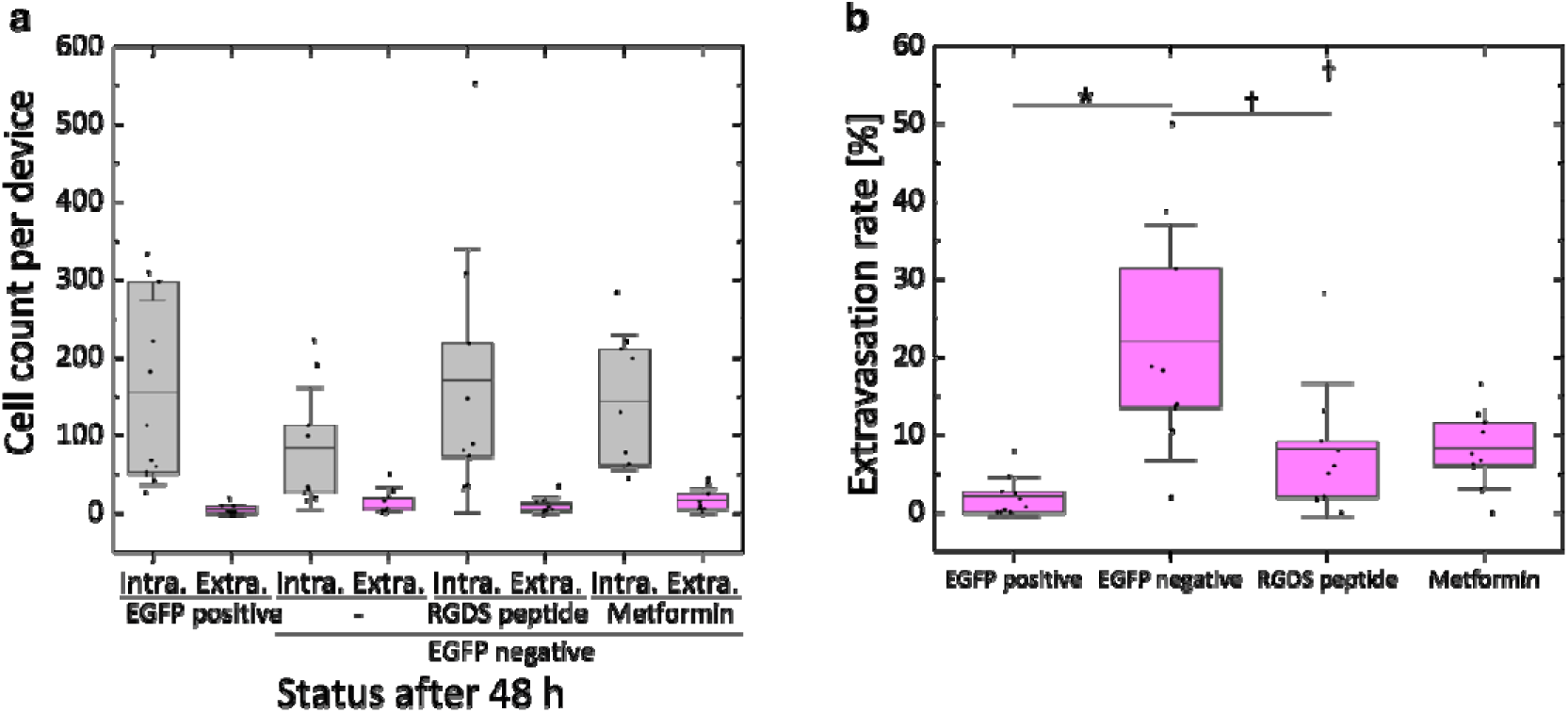
Comparison of (a)counts and (b)rate of extravasation of EGFP positive cells and EGFP negative cells with/without supplement of 0.5 M RGDS peptide (fibronectin/integrin inhibitor) or 48-h pretreatment of 5 mM Metformin (vimentin inhibitor). (*N* = 3, *n* ≥ 9). Error bars show standard deviation. Significant changes compared to EGFP negative without drug conditioning were assessed by *t*-test.

**Table. S1:**
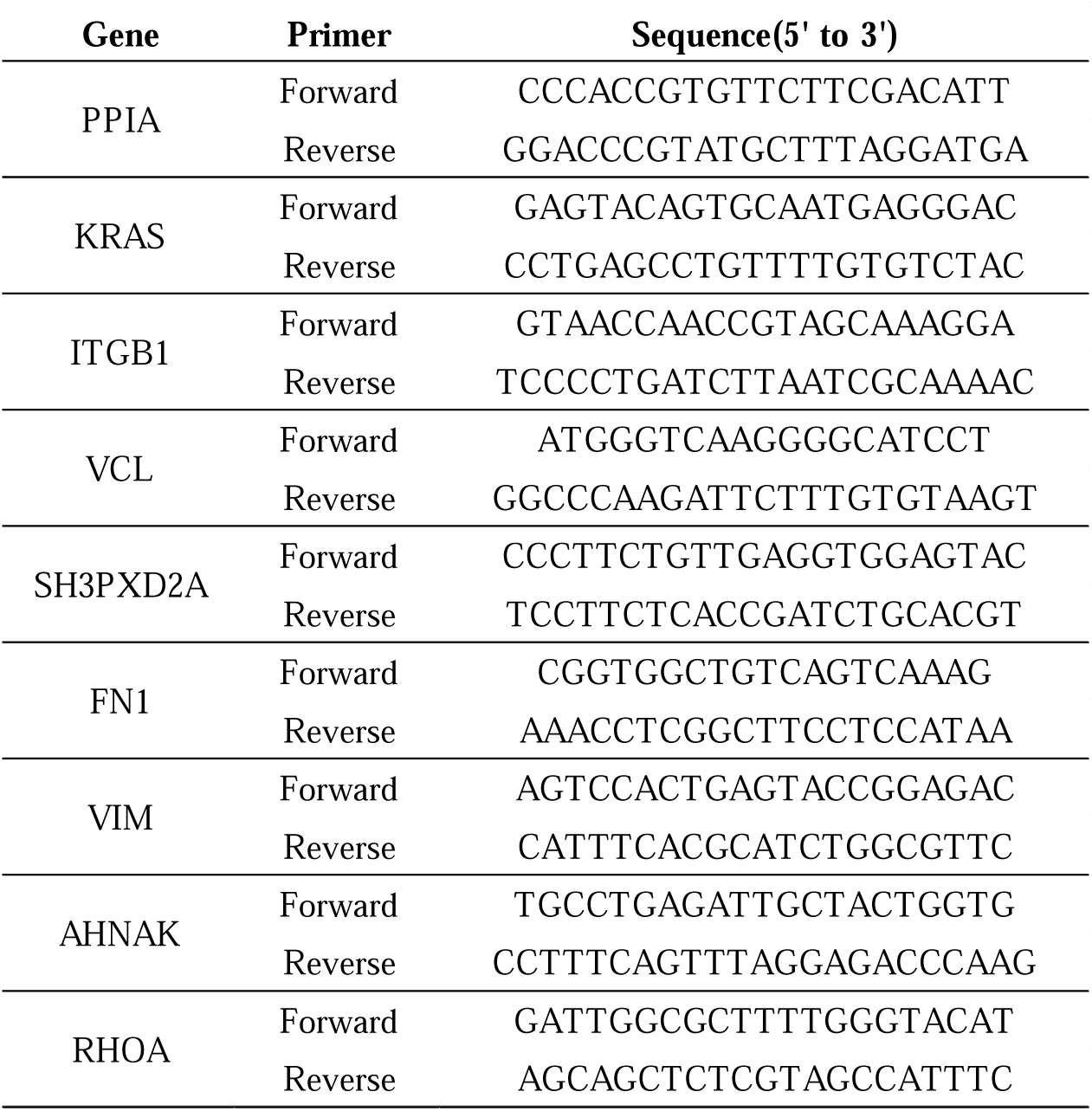
Primers used for qPCR.

